# Machine learned potentials with electrostatic embedding accurately capture Kemp eliminase reactivity

**DOI:** 10.64898/2026.09.14.751485

**Authors:** Abbie Lear, Elliot W. Chan, Kirill Zinovjev, Marc W. van der Kamp, H. Adrian Bunzel, Adrian J. Mulholland

**Affiliations:** Centre for Computational Chemistry, School of Chemistry, University of Bristol, Bristol, UK; Departamento de Química Física, Universidad de Valencia, 46100, Burjassot, Spain; School of Biochemistry and Biomedical Sciences, University of Bristol, Bristol, UK; Max Planck Institute for Terrestrial Microbiology, 35043 Marburg, Germany

## Abstract

Kemp elimination has become a benchmark for *de novo* enzyme design due to its simplicity and detailed mechanistic understanding but accurate barrier calculations are required to understand the difference in activity between designed Kemp eliminases. QM/MM MD simulations are capable of calculating reaction barriers, but are limited by a cost/accuracy tradeoff in which the most accurate methods are too computationally expensive for extensive screening as would be required in a prospective enzyme design campaign. Recent advances in embedded ML/MM simulations using the electrostatic machine learning embedding (EMLE) method have enabled transferable potentials for ML/MM MD simulations with low computational cost but QM-level accuracy. Here, we trained a MACE MLIP and EMLE embedding model for fast and accurate simulations of the enzymatic, base-catalysed Kemp elimination of 6-nitro benzisoxazole. Applying our EMLE ML/MM scheme to a designed Kemp eliminase and its evolved counterpart captured the *∼*4 kcal/mol difference in barrier between the two observed in experiment. Furthermore, training was done using structures generated from simulations of a single variant and was applied to the second variant, achieving near-experimental barriers, with no further finetuning or computational overhead. Thus, this work establishes EMLE-based ML/MM MD simulations as a potential route for fast and accurate assessment of barriers in computational screening for enzyme design.

---

Many enzymes are highly active and selective catalysts which operate under mild conditions and so have become increasingly useful in ‘green’ chemical synthesis.^1^ Computational *de novo* enzyme design, driven by machine learning models, has shown success in creating biocatalysts for new-to-nature reactions.^2–7^ However, current design protocols usually fall short of the rates seen in natural enzymes and designed enzymes can often be further improved by directed evolution.^8–13^ Understanding how evolution changes enzyme activity could shed light on factors that contribute to enzyme activity. Accurately modelling enzyme catalysis with simulations to assess reaction barriers and link these to structural effects could unlock new levels of computational enzyme design.

Quantum mechanics/molecular mechanics (QM/MM) molecular dynamics (MD) simulations have provided significant insight into the function and mechanisms behind many enzyme-catalysed reactions, but are currently limited in scope.^14^ Using density functional theory (DFT) to describe the QM region allows for accurate modelling of chemical reactions, but limits the available sampling due to high computational cost, and thus cannot be used for screening many enzymes. Semi-empirical methods have been developed to address this but they often produce inaccurate energies and geometries when compared to higher levels of theory,^15^ but are significantly cheaper for running MD simulations. Recently, machine learning interatomic potentials (MLIPs) such as MACE,^16^ NequIP^17^ and ANI^18^ have been developed that offer near-DFT accuracy at a fraction of the cost.^19^ However, embedding MLIPs into MM environments remains difficult, particularly as many reactive MLIPs are trained to be system specific and do not account for surrounding environments. This is particularly important for enzyme catalysis, as the electrostatics of the surrounding protein often polarises the reactants and transition state to a different extent, affecting the activation free energy and therefore the reaction rate.

The electrostatic machine learning embedding (EMLE) framework is one way to address this, predicting electron density and atomic polarisabilities of the reactive region to capture its response to the surrounding environment.^20–22^ Because the EMLE framework is agnostic to the MM environment, once the MLIP and EMLE embedding model are trained on a specific ML region, they can be applied in a variety of enzyme environments. Furthermore, both EMLE and MACE models are highly data efficient, requiring only a few hundred to thousand structures to be able to accurately predict energies, significantly reducing the required amount of training data. This fast and accurate method for modelling reaction barriers facilitates computational screening of enzyme variants, and has previously been able to separate across a class of enzymes by reaction barriers with no retraining.^23^

Kemp elimination (Fig. 1a) has become a benchmark reaction for *de novo* enzyme design.^4,5,24–26^ 1A53-2 is a computationally designed Kemp eliminase, but it has limited activity (*k*_*cat*_ = 0.0058 s^*−*1^).^25^ 1A53-2 was improved by directed evolution, which allowed boosting activity by more than 3 orders of magnitude with only six mutations. (*k*_*cat*_ = 10 s^*−*1^) with only six point mutations (Fig. 1b,c).^8^ Umbrella sampling simulations using the EMLE-based ML/MM reproduced reaction barriers within 1 kcal/mol of experimental data for both variants. Thus, we show that training models for a single system can successfully predict experimentally derived ΔΔ*G*^*‡*^ between the variants without retraining, suggesting a practical route to computationally guided enzyme design.

**Figure 1.**
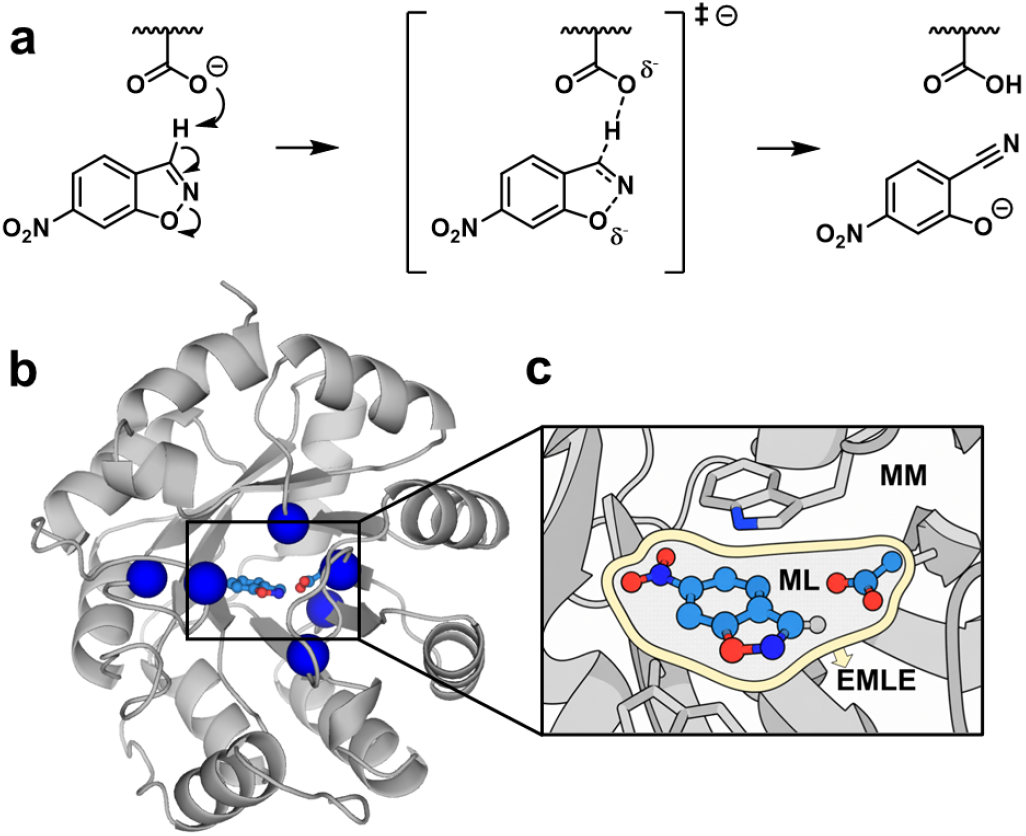
a) The base-catalysed Kemp elimination. b) The structure of 1A53-2 with the mutations introduced by directed evolution shown in blue spheres.^8^ c) The EMLE scheme: the ML region is treated using an MLIP and the EMLE model reads the point charges of the MM region and predicts how the ML region would respond. The MM region is treated normally.

To train the EMLE and MACE models, a query-by-committee active learning protocol was used (Fig. 2a). Initial structures for the first round of training were extracted from previous PM6/CHARMM36 reaction barrier calculations of the Kemp elimination of 6-nitro-benzisoxazole in 1A53-2 using the adaptive string method (ASM).^27,28^ Three MACE MLIPs and a single EMLE model (QbC is only applied to gas-phase MACE forces) were trained on energies and forces of the isolated QM region from single-point calculations performed using ORCA 6.0.1^29^ at the M06-2X^30^/6-31G(d) level. Umbrella sampling (US) ML/MM dynamics were then run using the trained MLIP and EMLE models. The collective variable (CV) was defined as a linear combination of three distances (LCod) found using the ASM reaction path. At every step of the simulation, all three MLIPs predicted energies and forces to check committee diversity. Structures outside of the training set were identified by a force deviation greater than 0.005 Hartree/Bohr between the committee of models. Representative structures with high deviation were added into the training set and the MACE and EMLE models were retrained. After three rounds of active learning the deviation between models across the reaction coordinate was significantly reduced (Fig. 2b) and the resulting MACE potential and EMLE model showed good fits to the training and test datasets (Fig. S1).

**Figure 2.**
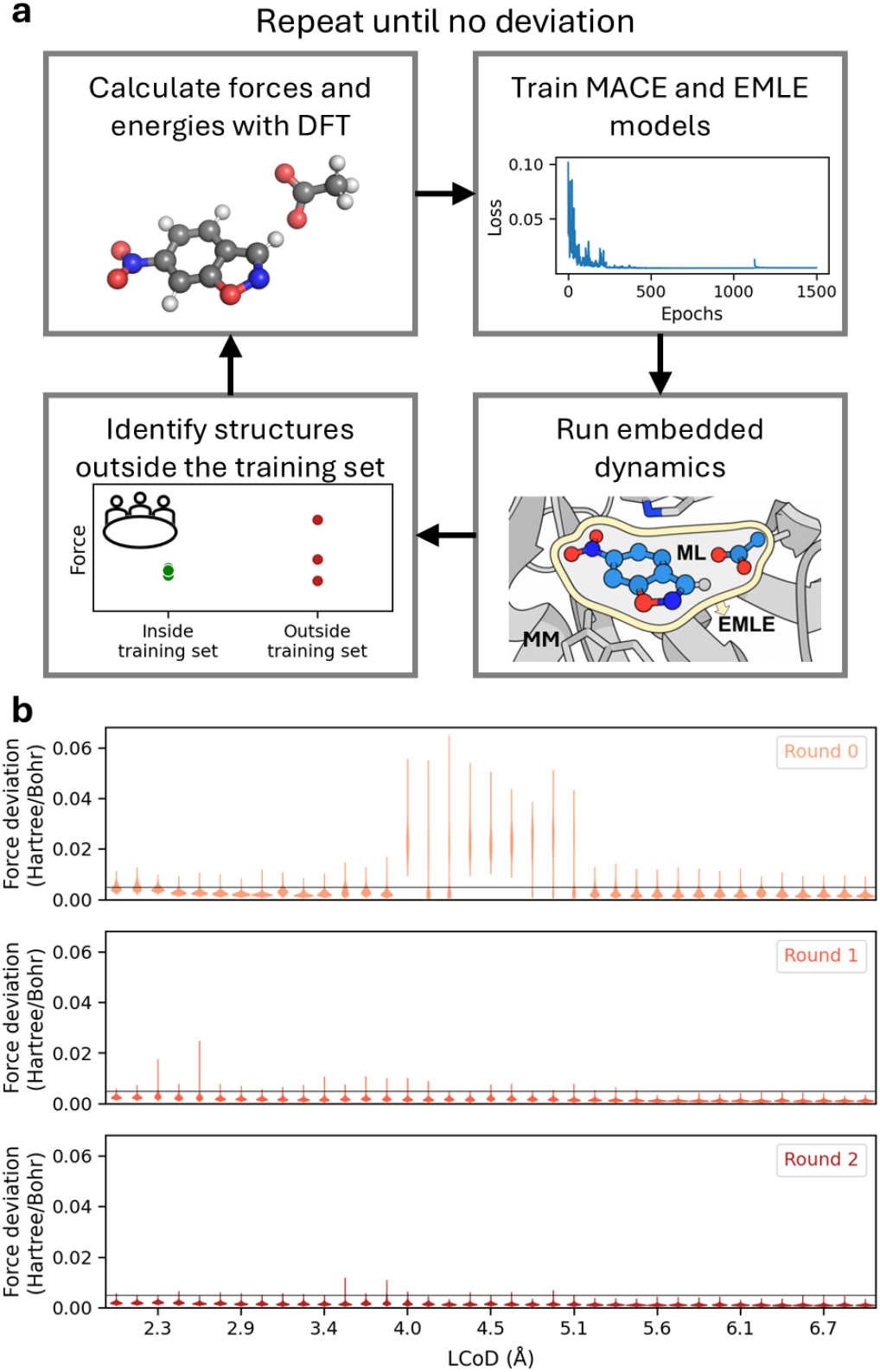
a) The active learning protocol used to develop the training set for the MACE and EMLE models used in this work. b) Deviations across a committee of three models identified structures to add for active learning. The cutoff applied was 0.005 Hartree/Bohr as marked on the figure. See supporting information for detailed methods.

Using the trained models, five independent umbrella sampling simulations were performed for both 1A53-2 and 1A53-2.5 in order to calculate the average barrier to reaction (Fig. 3). Starting structures were taken from previous MD simulations. These runs were initiated from their starting structures at a reaction coordinate value of 3 and propagated in both directions using the endpoint of the previous window to start the next.

**Figure 3.**
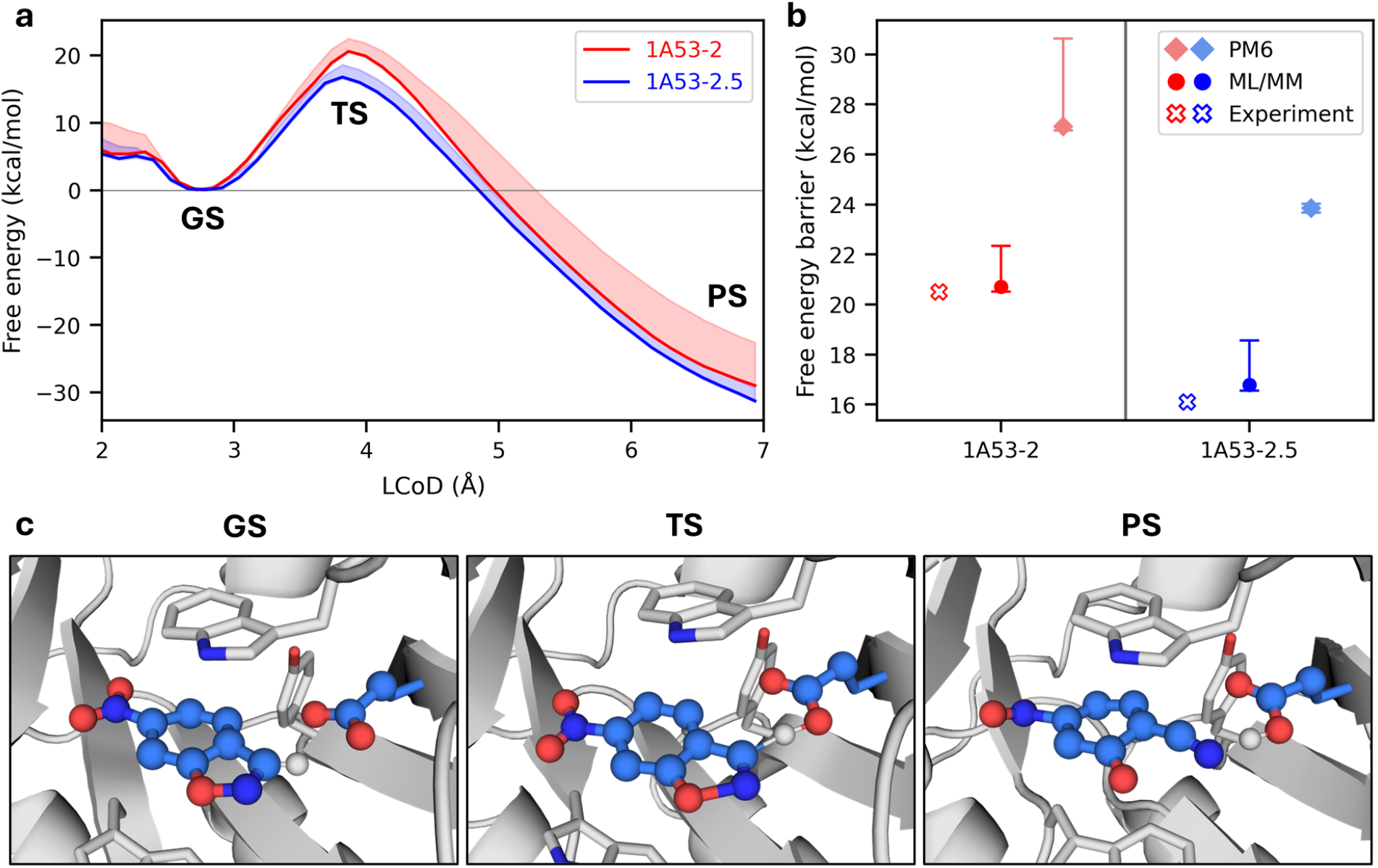
a) Potential of mean force from umbrella sampling of 1A53-2 and 1A53-2.5 (ML/MM with EMLE, solid lines show Boltzmann average over 5 repeats and shaded areas are the 95% confidence interval). b) Average barrier from PM6/MM and ML/MM with EMLE simulations compared to experiment (error bars are the 95% confidence interval). c) Representative ground state (GS), transition state (TS) and product state (PS) structures.

Visual inspection of the generated structures suggests that this MACE and EMLE methodology reproduces the expected chemistry (Fig. 3c). Calculating a Boltzmann average barrier for each gives 20.7 *±* 2.9 kcal/mol for 1A53-2 and 16.8 *±* 1.9 kcal/mol for 1A53-2.5 which gives a ΔΔ*G*^*‡*^ of 3.9 kcal/mol, consistent with the ΔΔ*G*^*‡*^ of 4.4 kcal/mol determined from the experimental kcat values using transition state theory.^8^ Furthermore, both calculated reaction barriers are within 1 kcal/mol of the experimentally derived barriers (1A53-2: 20.5 kcal/mol, 1A53-2.5: 16.1 kcal/mol). Such clear separation and capture of the relative change in barrier between the variants implies that further variants could be accurately screened and ranked computationally.

A variation of up to 7 kcal/mol is observed between replicates (Fig. S2), which is expected due to conformational changes that alter the reaction barrier. Effects like structural organization, solvation, and electrostatic arrangement at the active site can alter the barrier^27^ and a spread of barriers is also seen when using semi-empirial QM/MM simulations (PM6/ff14SB) with the adaptive string method (Fig. 3b, S2). To ensure these effects are accounted for, Boltzmann average barriers were calculated, such that the enzyme-substrate conformations that produce the lowest reaction barriers contribute most significantly to our calculated value. When using a Boltzmann average and a bootstrapped confidence interval, these ML/MM simulations achieve near-experimental accuracy with five replicates.

We analysed the performance of our EMLE and MACE models with reference to single point DFT(M06-2X/6-31G(d))/MM calculations using the emle-analyze tool (Fig. S3). The MACE model accurately captures the *in vacuo* energy of the reactive region compared to the DFT energies. The difference in EMLE embedding energy varies along the reaction coordinate, similar to previous ML(EMLE)/MM simulations for other systems.^22^

The computational efficiency of the ML/MM simulations was compared to equivalent DFT/MM and PM6/MM MD simulations by measuring how long is required to perform a single step of the simulation (in order to take the efficiency of the MD engine into account). The ML/MM simulations were *≈*500 times faster than DFT/MM, on the order of PM6/MM simulations (Table S1), whilst predicting significantly improved reaction barriers than the PM6/MM simulations. We note that we used the original emle-engine implementation for this test, which uses a separate emle-server process for the ML/MM calculations with sander.[21] Future implementations will not require the separation, substantially reducing file I/O. Preliminary tests of the serverless interface, performed on a separate cluster, show a step time *≈*3200 times faster than original DFT/MM (Table S2) using 16 CPU cores for the MM calculation and a GPU for the ML calculations. Also the ML region used here was quite small at only 22 atoms. MLIPs scale significantly better than either semi-empirical or DFT methods and so as ML region size increases the speed-up versus QM/MM will be more notable.

We have demonstrated the application of combined ML(EMLE)/MM simulations to compare two variants of the Kemp eliminase 1A53-2. Trained solely on geometries extracted from simulations of a single variant, the model accurately predicted experimental reaction barriers between variants, capturing the *≈* 4 kcal/mol enhancement of the evolved variant. This workflow represents a practical, high-throughput route towards computationally screening *de novo* enzyme designs and accurately determining their capabilities as biocatalysts.

## Author Contributions

AL performed the training and simulations with assistance from EWC and KZ and supervision from MWVDK, HAB and AJM. All authors contributed to data analysis and discussion. AL and EWC wrote the initial manuscript; all authors contributed to manuscript revisions.

## Data Availability

Data that support the findings of this study are available within the paper and its Supplementary Information. Simulation inputs and ML models will be made freely available upon manuscript acceptance.

## Acknowledgements

This work is part of a project that has received funding from the European Research Council under the European Horizon 2020 research and innovation programme (PREDACTED Advanced Grant Agreement no. 101021207) to AJM. HAB thanks the SNSF (PZ00P3_208691), the Max Planck Society, and Max Planck Foundation for support. EWC and MvdK acknowledge EPSRC grant EP/V011421/1 ‘Simulating catalysis: Multiscale embedding of machine learning potentials’. KZ acknowledges grant RYC2024-051397-I, funded by MICIU/AEI/10.13039/501100011033 and by FSE+. KZ also acknowledges Europa Excelencia grant EUR2025-165005 funded by Spanish State Research Agency (AEI). This work was conducted using the computational facilities of the Advanced Computing Research Centre, University of Bristol (http://www.bris.ac.uk/acrc/). We also thank Barcelona Supercomputing Center (project QH-2026-1-0004) for computational resources.

