## Supplementary Information for "Machine learned potentials with electrostatic embedding accurately capture Kemp eliminase reactivity"

#### S1 Supplementary Methods

##### S1.1 Adaptive String Method

String method reference simulations are taken from this work.[1] Briefly, 32 nodes and four collective variables detailed below (Table S3) were used in these simulations and starting conformations were taken from multi-replicate timescale molecular dynamics simulations (4 snapshots from 10 500 ns simulations). The minimum free energy path was optimised over 20 ps of simulation and then sampled over 100 ps of simulation using PM6/CHARMM36.

##### S1.2 Initial Structures

Initial structures for training the initial models prior to active learning were taken from the previous string method profiles. QM region structures were taken from the final frame at each string node from 38 successful string method profiles calculated for 1A53-2 with a transition state model of the ligand yielding a total of 1216 structures.[1] 1A53-2 was selected as showed higher structural diversity than 1A53-2.5 and the ensemble generated with the TS model of the ligand was selected as it showed better reactive behaviour than the GS model. Link atoms were added to the QM region structures using emle-server[2] to capture sander's[3] output.

##### S1.3 DFT calculations

All DFT calculations were performed with the M06-2X functional and the 6-31G(d) basis set using ORCA v6.0.1. The molecular polarisabilities were also calculated for each structure to faciliate EMLE training as well as saving the electronic density. MBIS decomposition for EMLE was performed with HORTON.[4]

#### S1.4 EMLE and MACE active learning protocol

A Query-by-Committee active learning approach similar to our previous work[5] was employed to train both the EMLE model and MACE MLIP. The initial training set contained 1216 structures. The transition state showed high deviation between models, indicating that this area of conformational space was not well represented by the training set. Up to five structures from each window, which deviated between 0.005 and 0.02 Hartrees/Bohr in force prediction, were selected to be added to the training set, yielding a total of 247 extra structures to give a total training set size of 1464 structures. In the second round of active learning the model performed better again, with only four windows showing significant deviation and 126 structures were added to give a total training set of just 1591 structures.

1. Initial structures were taken from the string method calculations (see above)
2. Single point calculations were performed at the M06-2x/6-31G(d) level as described above.
3. The EMLE model was trained using the emle-engine package.[2] The default parameters were used except for increasing the number of epochs to 200 and a threshold of 0.02 for invariate vector machine used to select basis set atoms for learning valence widths and electronegativities.
4. Each MACE potential used an equivariant message-passing architecture with two interaction layers, hidden features carrying irreducible representations up to L=1 across 128 channels, and a radial cutoff of 5.0 Å. The reference dataset was partitioned into training and test sets in an 80:20 ratio, with a further 5% of the training partition held out for validation during fitting. Atomic reference energies (E0s) were taken from isolated-atom reference calculations rather than regressed from the dataset. Models were fitted to the DFT energies and forces using the Adam optimiser with a batch size of 10 for a maximum of 1500 epochs, and stochastic weight averaging was applied in a second training stage to refine the final parameters. Committee diversity was obtained by training 3 potentials that differed in their random initialisation seed.
5. Dynamics were performed using the trained EMLE and a single of the MACE MLIPs. At every step, all 3 MACE MLIPs predicted energies and forces for the ML region, with each prediction being recorded to be used as the metric for model uncertainty.
6. Structures which deviated in force prediction between 0.005 and 0.02 Hartree/Bohr were considered to be poorly represented in the current training set. Up to 5 structures were selected per window, a minimum of 100 MD steps apart, and added to the training set.
7. The models were retrained
8. Repeated until all windows were within an acceptable tolerance. The final model's fit to training and test sets can be found in Figure S1.

#### S1.5 ML/MM simulations

All dynamics were performed using AMBER24[3] using the EMLE server[2] that intercepts AMBER's call to an external QM program. The ML region consisted of the ligand and catalytic base from C$\gamma$ onwards so that the C$\beta$-C$\gamma$ bond crossed the boundary in a similar protocol as described in.[1] The cutoff used for MM atoms included in the EMLE calculations was 16 Å (consistent with QM/MM implementations using external interfaces in sander from AmberTools24). A 1 fs timestep was used for all simulations. Starting structures and simulation parameters were taken from a previous study.[1]

The reaction coordinate used was a linear combination of distances (LCOD) combining the scissile C-H bond distance, the ring opening N-O bond distance and the forming H-C$\delta$ bond distance. These distances were scaled by 0.5, 2 and -0.2 respectively to mimic the rates of change seen in ASM. This protocol was first tested using PM6/MM from an MD generated start point. For each US window, 2 ps of equilibration was performed before 10 ps of production. Initial structures for the ML/MM runs in training were taken from the PM6/MM endpoints at each window. Each window was spaced 0.1 Å apart, resulting in 51 windows (from 2.0 to 7.0). Production runs using the trained models were run in a sequential manner, starting from RC = 3.0 and proceeding forwards and backwards, using the previous window's endpoint to start the next. For each restraint, a force constant of 200 kcal/mol was applied, and the Grossfield implementation of the WHAM was used to integrate the PMF.

#### S2 Supplementary Figures

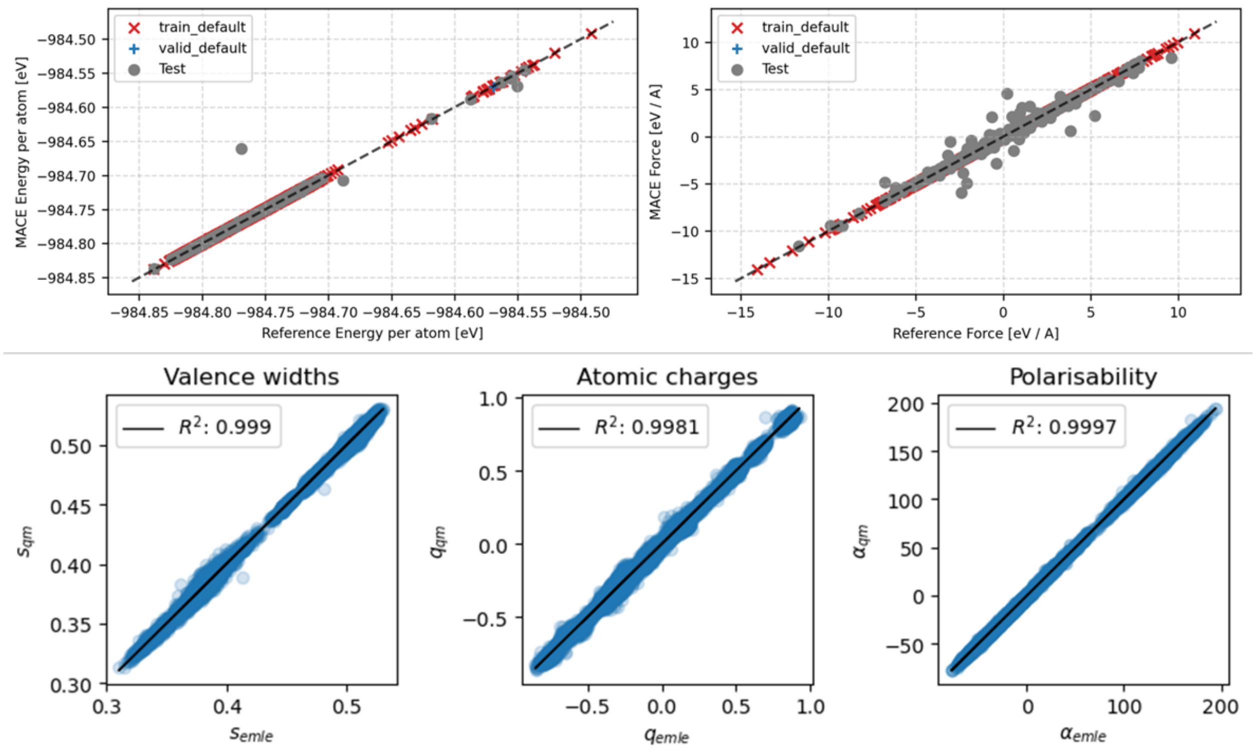


Figure S1: MACE and EMLE training results showing a good fit to the data. The top two panels show the correlation of MACE MLIP energies and forces to the reference DFT. The bottom three panels show the correlation between the parameters learnt by EMLE for electrostatic fitting and the reference values from DFT.

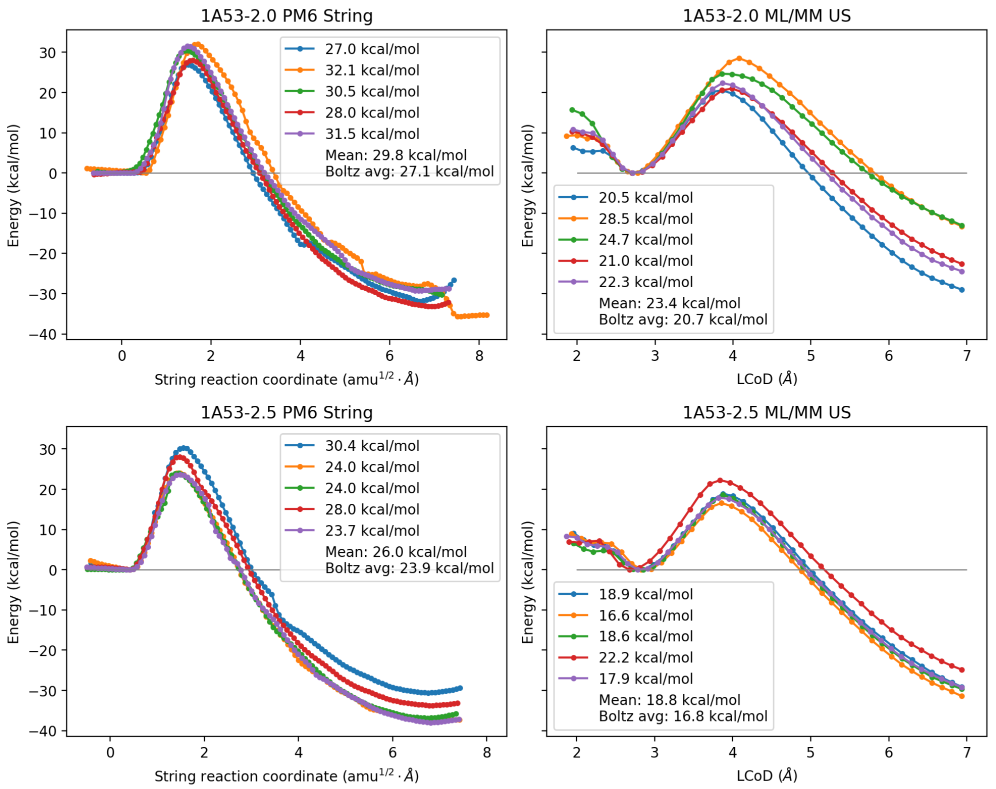


Figure S2: Comparison of PM6 reaction profiles (calculated with the adaptive string method) with ML/MM (calculated with umbrella sampling).

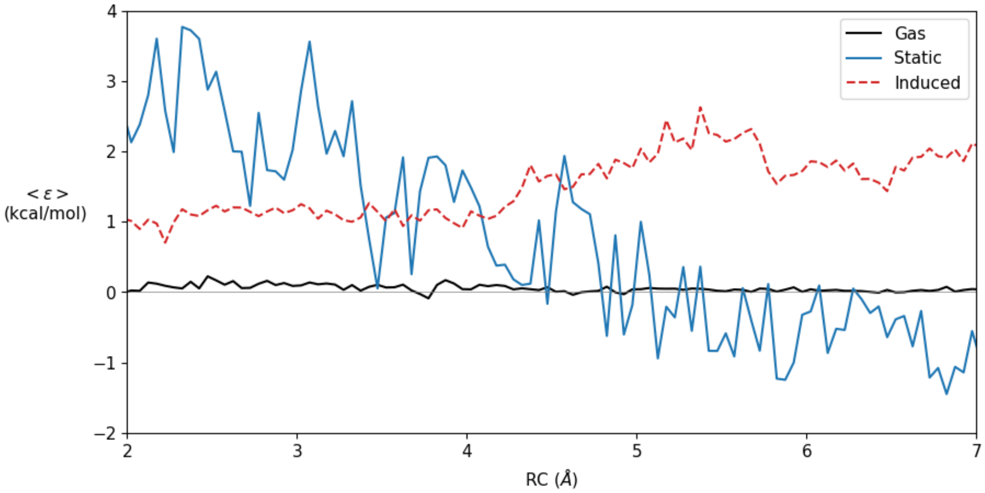


Figure S3: Comparison of EMLE embedding to QM electrostatic embedding using the emle-analyze function. The error is broken down into three components: the error in the gas-phase MLIP energies (black), the error in the interaction energy of the static electronic density of the ML region with the surrounding MM region (blue) and the error in the energy of forming the induced dipoles on the ML region and their interaction with the MM region (red, dashed)

### S3 Supplementary Tables

Table S1: Comparison of sampling speeds between different methods (single core for QM(ML) and MM calculation, except for one example offloading the EMLE server and MLIP onto a GPU). All calculations were performed on an Intel Xeon w7-3445 CPU, with the GPU calculation performed on a NVIDIA RTX 4500 Ada GPU using emle-server interfaced with AMBER24.

| **Method** | **Time (ms/step)** |
|---|---:|
| M06-2X | 130000 |
| PM6 | 140 |
| $EMLE_{CPU}$ | 260 |
| $EMLE_{GPU}$ | 200 |

Table S2: Sampling speeds when using the serverless interface between sander and EMLE, with a varying number of cores to perform the MM calculations whilst the ML calculations are performed on the GPU. Performed on MareNostrum 5 with Intel Xeon Platinum 8480+ CPUs and Nvidia H100s.

| **CPU cores** | **Time (ms/step)** |
|:---:|:---:|
| 1 | 130 |
| 2 | 86 |
| 4 | 59 |
| 8 | 47 |
| 16 | 41 |
| 20 | 47 |

Table S3: Collective variables used for the adaptive string method reaction barrier calculations.

| **Collective variable** | **Start value** | **End value** |
|---|---|---|
| C – H distance | 1.07 Å | 3.20 Å |
| N – O distance | 1.43 Å | 3.60 Å |
| C – C – N angle | 110° | 180° |
| H – C$\delta$ distance | 2.80 Å | 2.00 Å |
